# Macrophage plasticity is Rac signalling and MMP9 dependant

**DOI:** 10.1101/614388

**Authors:** Jana Travnickova, Sandra Nhim, Naoill Abdellaoui, Farida Djouad, Maï Nguyen-Chi, Andrea Parmeggiani, Karima Kissa

## Abstract

*In vitro*, depending on extracellular matrix (ECM) architecture, macrophages migrate either in amoeboid or mesenchymal mode; while the first is a general trait of leukocytes, the latter is associated with tissue remodelling via Matrix Metalloproteinases (MMPs). To assess whether these stereotyped migrations could be also observed in a physiological context, we used the zebrafish embryo and monitored macrophage morphology, behaviour and capacity to mobilisation haematopoietic stem/progenitor cells (HSPCs), as a final functional readout. Morphometric analysis identified 4 different cell shapes. Live imaging revealed that macrophages successively adopt all four shapes as they migrate through ECM. Treatment with inhibitors of MMPs or Rac GTPase to abolish mesenchymal migration, suppresses both ECM degradation and HSPC mobilisation while differently affecting macrophage behaviour. This study depicts real time macrophage behaviour in a physiological context and reveals extreme reactivity of these cells constantly adapting and switching migratory shapes to achieve HSPCs proper mobilisation.

## Introduction

Macrophages were for the first time identified as phagocytic cells responsible for pathogen elimination (Metchnikoff, 1892). Over the past century, they were associated with homeostasis, innate and adaptive immune responses, inflammation, tissue remodelling and cytokine production (Gordon and Taylor, 2005; Wynn et al., 2013). Macrophages are the most plastic haematopoietic cells present in all tissues; their diversity depends upon their location, their morphology, their membrane receptors or surface markers (Wynn et al., 2013). Depending on tissue composition they infiltrate and environmental constraints, macrophages adopt different migration modes (Vérollet et al., 2011). In the course of a three-dimensional (3D) migration, macrophages can either adopt an amoeboid or a mesenchymal migratory mode. In case of an amoeboid migration, cells take on a round or polarised shape and migrate through the extracellular matrix (ECM). Such a migration is Rho/Rock GTPases dependent. During mesenchymal migration, macrophages degrade the ECM through proteinases secretions (e.g. Matrix Metalloproteinases or MMPs) and cells take on an elongated shape. This second migratory mode is Rac GTPase signalling dependent (Sanz-Moreno and Marshall, 2010; Vérollet et al., 2011).

In mouse and human, macrophage characterization was mainly performed *in vitro* using bone marrow derived macrophages. Recently, the zebrafish model was used to resolve specific issues during the developmental process or to address accurate pathologies. The transparency of zebrafish embryos enables the live imaging and real time tracking of cell populations. We and other groups have shown that two main waves of macrophages emerge from primitive and definitive haematopoiesis during the zebrafish development (Gering and Patient, 2005; Herbomel et al., 1999; Murayama et al., 2006). The initial wave takes place between 18 and 25 hours post fertilization (hpf) in the yolk sac (Herbomel et al., 1999). The second wave occurs between 30 and 55 hpf in the aorta-gonad-mesonephros (AGM) (Gering and Patient, 2005; Murayama et al., 2006) and generates the haematopoietic stem/progenitor cells (HSPCs) (Bertrand et al., 2010; Kissa and Herbomel, 2010) which later will differentiate into all blood cells including macrophages. Finally, a transient hematopoietic wave is initiated in the posterior blood island, giving rise to the multilineage progenitor cells and erythromyeloid progenitors, which develop into both erythroid and myeloid cells (Bertrand et al., 2007).

Recently, we demonstrated *in vivo* that primitive macrophages are crucial in the establishment of a definitive haematopoiesis (Travnickova et al., 2015). Macrophages that accumulated in the AGM degrade the ECM located in the vicinity of HSPCs via matrix metalloproteinase 9 (MMP-9) secretions, thereby enabling them to migrate, enter the blood stream and colonise haematopoietic organs.

In the present study, we provide an extensive analysis of macrophages in zebrafish embryos. Using morphological analysis we were able to distinguish for the first time different macrophage subtypes *in vivo*. By combining morphological analysis with live imaging we succeeded in visualizing the dynamic behavioural patterns of individual macrophages during their migration through the ECM.

## Results

### Macrophage shape heterogeneity in the zebrafish embryo

During the establishment of the definitive haematopoiesis, macrophages accumulated in the AGM between 30 and 60 hpf degrade the ECM surrounding HSPCs via Mmp-9 secretions which result in the mobilization of HSPCs (Travnickova et al., 2015). Using this physiological model, we analysed the shape and behaviour of proteolytic macrophage in order to establish a potential correlation. We first described the position and shape of macrophages in the AGM using the *kdrl:eGFP//mpeg1:mCherry-F* double transgenic lines where the GFP protein highlighted vessels and mCherry-F macrophage membranes (**Fig. 1A-B**). Figure 1A provides a schematic view of the vessel and macrophage position as shown in Figure 1B. Using a 3D view (**Fig. 1B**) we were able to determine the position of macrophages (white arrows) in the outer layer of the vein wall between the vein and the aorta floor with different morphologies. The particle analysis of macrophages from a maximum projected confocal acquisitions enabled us to distinguish and quantify the various macrophage shapes. Three main morphological criteria were identified: circularity, roundness and elongation factor (**Fig. 1C and Suppl. Table 1**). They revealed the existence of 4 main shapes whose images are shown in **Figure 1D**. We named these 4 subgroups - round (1), amoeboid (2), star-like (3) and elongated shape (4). While the round and elongated shapes had already been described *in vitro*, the two remaining shapes might represent either subgroups present *in vivo* or intermediate stages between round and elongated shapes. The main difference between the amoeboid and star-like shapes lied in the presence in the amoeboid shape of a main axis, i.e. polarity. The quantification of each shape revealed that amoeboid, star-like and elongated shapes were equally present whereas the round shape remained sparse (**Fig. 1E**).

**Figure 1:**
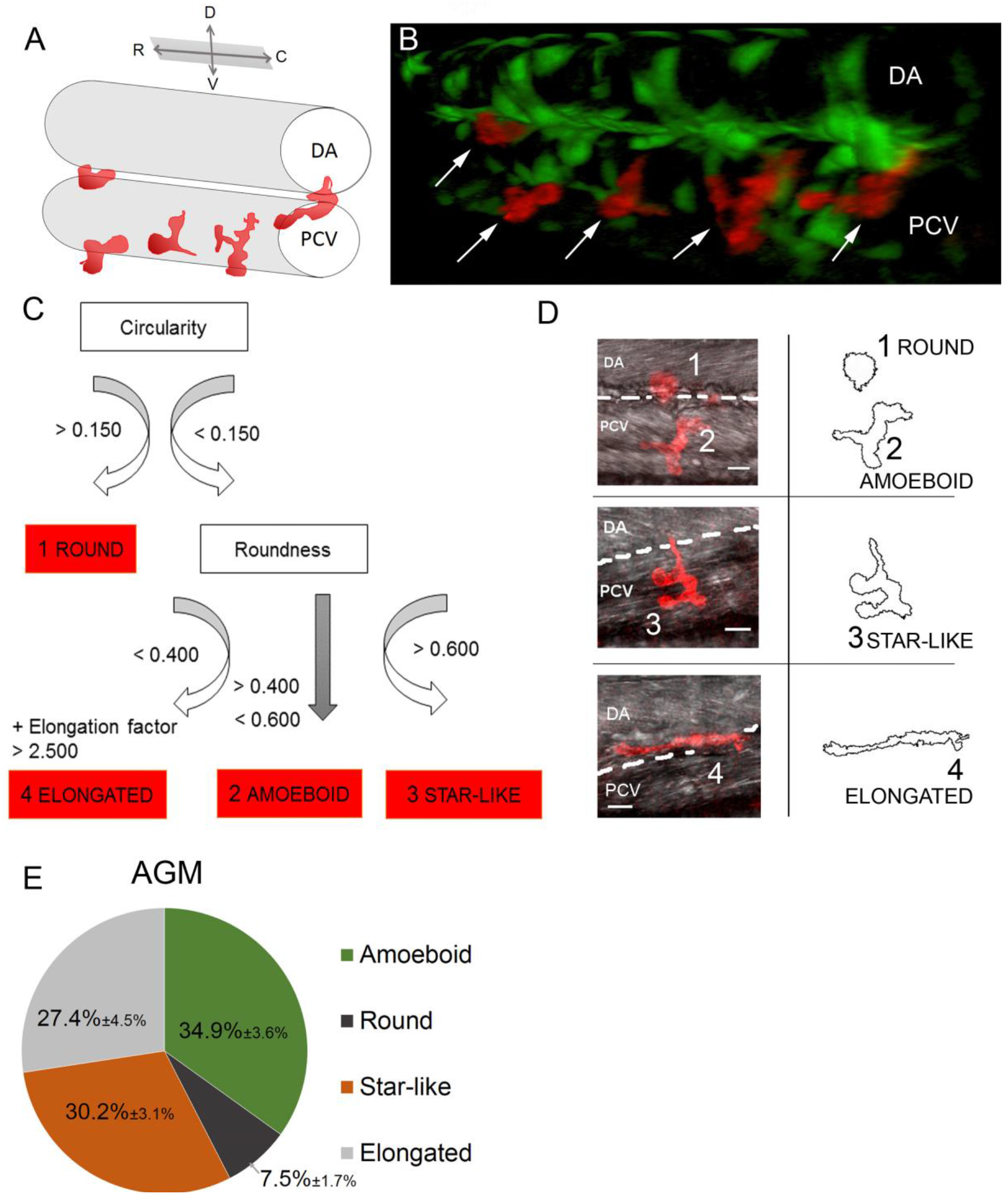
Macrophages in the AGM can be divided into 4 morphological subgroups. (A) The drawing shows a 3D view of vessels and macrophages (red) imaged in B. (B) 3D view of the AGM in the mid-trunk region of a *Tg(kdrl:eGFP//mpeg1:mCherry-F)* zebrafish embryo at 48 hpf showing the position of vessels (endothelium in green) and macrophages (in red, white arrows) in the outer side of the vein and in between the dorsal aorta and the cardinal vein. (C) Diagram of the 4 categories of macrophages delineated according to shape attributes–circularity, roundness and elongation factor. (D) Representative confocal images (maximum intensity projections) of individual categories with an outline drawing from particle analysis on the right. (E) Graph representing the percentage distribution of the different shape categories per AGM. Data are represented as percentage average ± s.e.m. N= 20 embryos. C, caudal; D, dorsal; DA, dorsal aorta; PCV, posterior cardinal vein; R, rostral; V, ventral. See also supplementary table 1.

### Dynamics of macrophage migration *in vivo*

The analysis of macrophage shapes revealed the existence of four morphological subgroups distributed in the zebrafish AGM. To assess the behaviour of each macrophage subgroup, we imaged *Tg(Mpeg1:mCherry)* embryos over the course of one hour (acquisition every minute; **Video 1.**). We selected time frames in colour depth projection that illustrated the dynamics of macrophages able to adopt different shapes within fifteen minutes (**Fig. 2A-F and Video 1,** colour code scale). The outlines represent the shape of macrophages in the imaged area at the 9th minute and enable us to draw a direct comparison with following time points. The colour depth projection of confocal imaging enabled us to determine the depth of macrophage positions *in vivo* and to demonstrate their ability to migrate in 3D patterns (**Video 1**). *In vivo* tracking of all macrophages within a 60 minute timeframe demonstrated that no specific directionality was maintained during their migration (**Fig. 2G**, n=23) as opposed to macrophages attracted to a wound site as an example of typical oriented migration (**Fig. 2H**, n=27). The speed of migration remained the same in both cases (data not shown). Subsequently, we quantified the evolution of macrophage shapes over time. Every single macrophage in the AGM reveals an ability to change shape within a very short time span (measured every 5 minutes) and to pass repeatedly through distinct shape subgroups over a 30 minutes course (**Fig. 2I**, n=10). The round shape appeared less frequently than others and live imaging showed that cells often adopted a round shape under two specific conditions: during cell division or once the macrophage entered the bloodstream.

**Figure 2:**
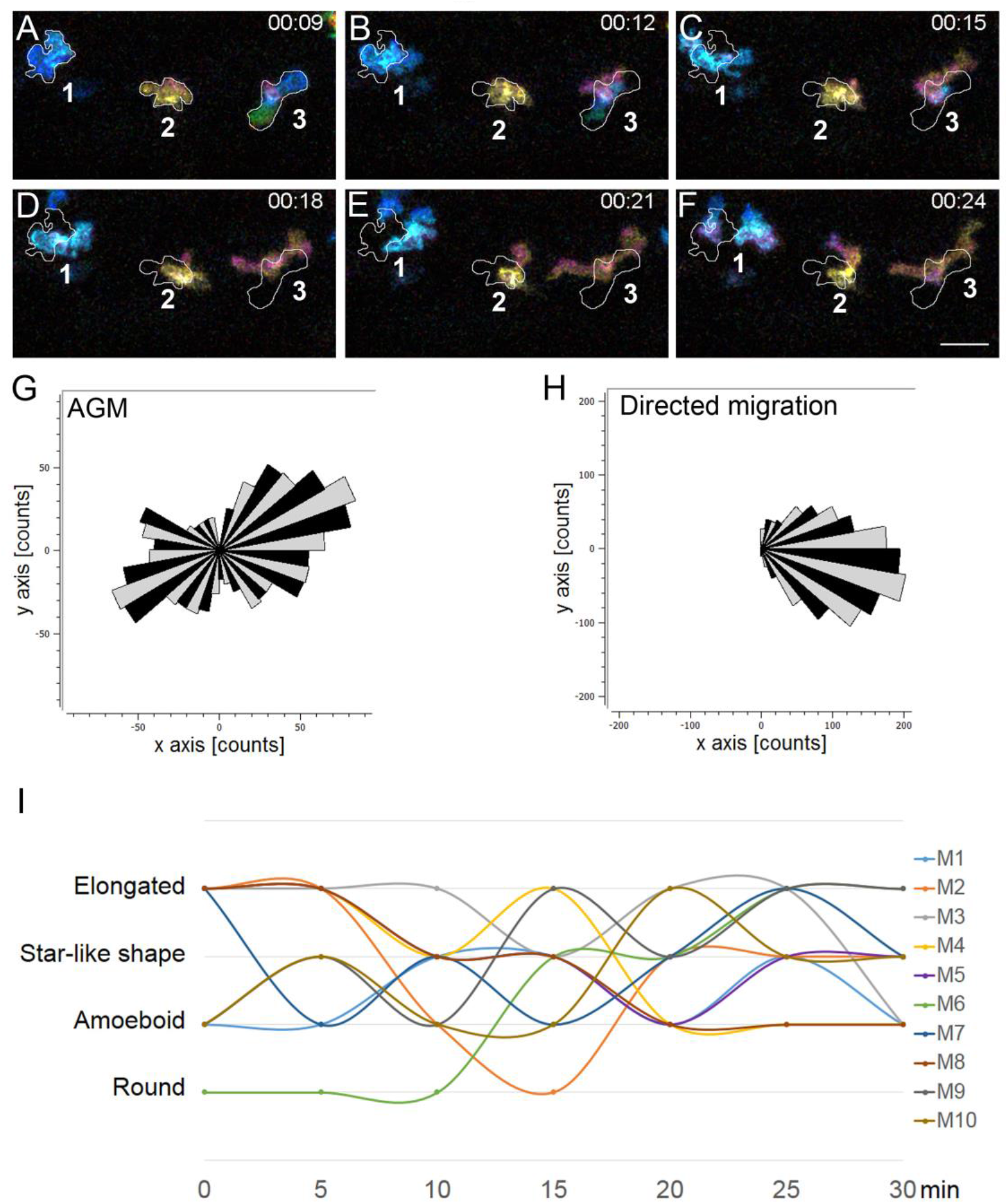
Macrophages in the AGM migrate in the mesenchymal way and undergo dynamic transition between different shapes over time. (A-F) Selected images from Video 1 illustrate the macrophage migration and shape transformation over time. Numbers point to individual macrophages. Time code is expressed in hours and minutes. White outlines on panel B-F indicate the shape and position of macrophages from panel A (9^th^ minute). (G-H) Rose plot diagrams show the directionality of macrophage migration in the AGM compared to the oriented migration of macrophages in the tail region after tail fin cut injury. A diagram represents the single counts of the position of each macrophage in the selected area (black and grey sectors of angle π/18) every minute over 60 minutes with a (x,y 0,0) starting point. n= 23 macrophages for the control and n=27 for directed migration. (I) Graph showing the shape evolution of individual macrophages during a 30 minutes course with 5 minutes interval measurements. Every line represents a single macrophage (n= 10). See also Video 1. Scale bar, (A-F) 30 μm.

In conclusion, macrophage real time imaging completes the characterisation of mesenchymal migrating macrophages and shows for the first time that they can adopt successive morphologies for their migration in the 3D matrix.

### Rac inhibition modifies macrophage behaviour and function

Using *in vivo* imaging we showed that macrophages exhibited morphological plasticity during their migration. This high plasticity depended on both, external (the stroma rigidity) and intrinsic parameters (cytoskeleton dynamics) (Vérollet et al., 2011). One intrinsic factor associated with mesenchymal migration is the small GTPase-Rac signalling. We thus investigated the effect of Rac chemical inhibition on macrophage shape and migration patterns. The macrophage shape distribution in Rac inhibitor (NSC23766) treated embryos did not significantly differ from that of DMSO treated control (**Fig. 3A**, N_DMSO_=10 and N_Rac inh._=15 embryos). Selected images from **Video 2** (colour, depth, projection, bottom) demonstrated that the macrophage migration was much slower than that of control embryos (**Fig. 3B-E, Video 2**, top). Macrophage speed measured over 60 minutes in the AGM confirmed a decrease in velocity from 2.37 ± 0.13 μm.min^−1^ to 1.13 ± 0.16 μm.min^−1^ (**Fig. 3F**).

**Figure 3:**
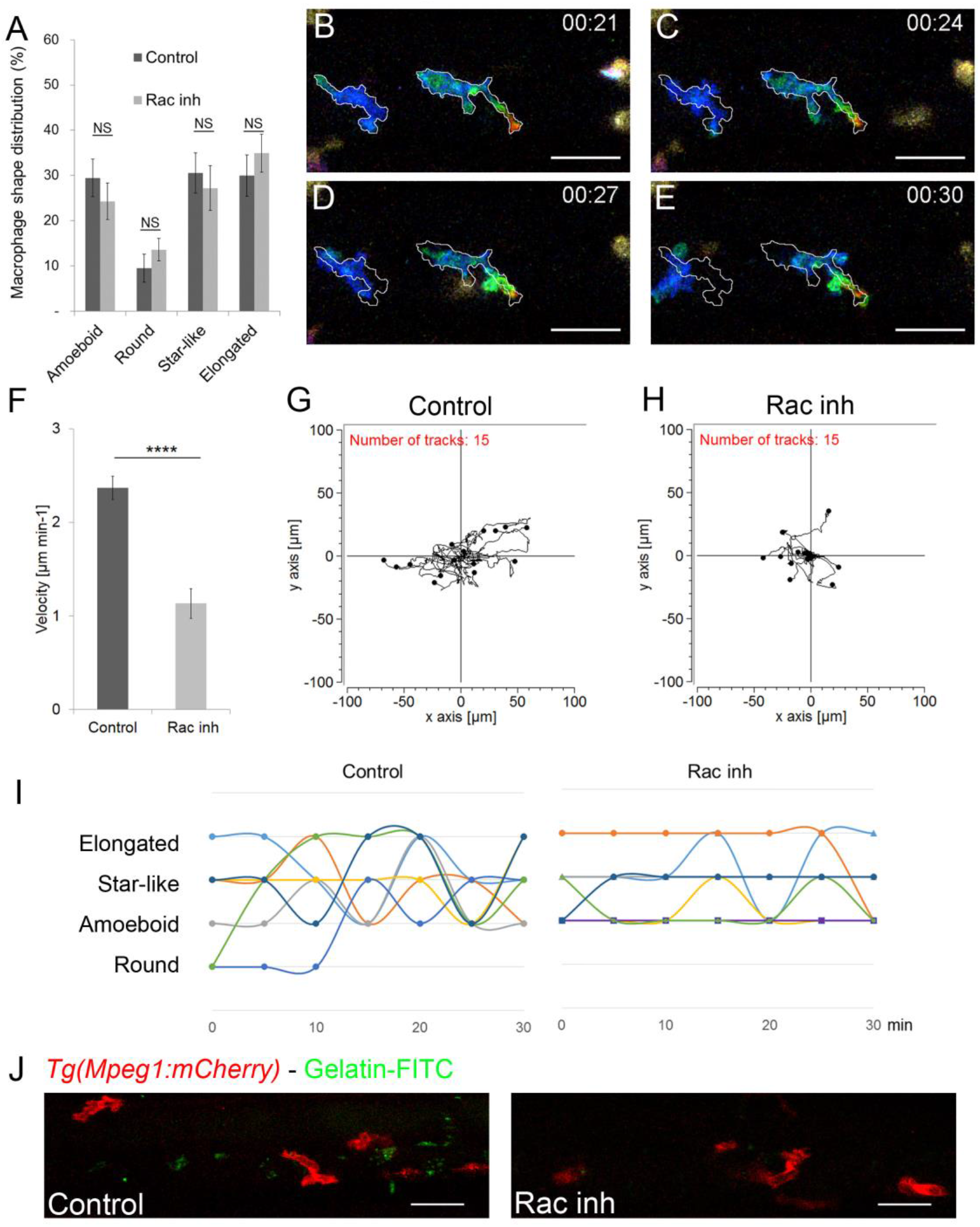
Rac inhibition leads to a loss of macrophage plasticity and motility. (A) Graph comparing macrophage shape distribution in the AGM of NSC23766 Rac-inhibited embryos (Rac Inh) to DMSO treated embryos (control) shows no significant change of distribution. N=10 embryos for control and 15 for Rac inh. Data are represented as the mean of the percentage of each shape type in the total macrophage population in the AGM ± s.e.m. NS = not significant. (B-E) Selected cropped images from Video 2 showing the shape and migration of macrophages over time. Time code in hours and minutes. White outlines on panel C-E indicate the shape and position of macrophages from panel B (21^st^ minute) (F) Graph showing the velocity of macrophages in control and Rac-inhibited embryos. Data are represented as a mean ± s.e.m., n= 15 macrophages from 4 different embryos, ****p<0.0001. (G-H) Tracking plot diagram representing the migration path and distance of macrophages in the AGM in control and Rac-inhibited embryos measured every minute for 60 minutes. Scale in μm, n=15 macrophages from 4 different embryos. (I) Graph shows the shape evolution of individual macrophages during a 30 minute course with 5 minutes interval measurements, Control to the left, Rac inhibitor to the right. Each line represents a single macrophage (n= 7). Statistically significant differences exist in the number of shapes adopted during a 30 minute measurement course (P=0.003) as well as in the number of changes between two different shapes (P=0.006). (J) *In vivo* zymography in *Tg(Mpeg1:mCherry)* embryos at 48 hpf reveals the degradation of inserted gelatin (green dots of cleavage-revealed FITC) in control embryos and a highly reduced degradation after Rac inhibition. See also Video 2. Scale bar: 30 μm.

The tracking plot diagram illustrated macrophage migration path and distance in control and Rac-inhibited embryos (**Fig. 3G-H**, n=15 macrophages from 4 embryos, position measured every minute over 1 hour) and revealed that Rac inhibition reduced macrophage moves from 130.9 ±7.5 μm to 56.8 ± 7.8 μm. Moreover, the analysis of macrophage shape dynamics, revealed a reduction in macrophage plasticity over time as macrophages were no longer able to consecutively adopt different shapes (**Fig. 3I, right**; n= 7) versus control (**Fig. 3I, left**; n= 7). However, in spite of reduced plasticity levels, membrane extensions were still formed at the same rate and with similar length as in control macrophages. Rac inhibition resulted in an increase in single extension span (from 2.8 ± 0.3 min to 11.0 ± 1.9 min, n_extension_= 45) as opposed to that of control macrophages.

As macrophage migration and morphological plasticity were significantly affected by Rac inhibition, we decided to evaluate the functionality of these macrophages. The main role of AGM macrophages is to degrade the ECM and to enable HSPC migration (Travnickova et al., 2015). *In vivo* zymography of *Tg(Mpeg1:mCherry)* embryos at 48 hpf, revealed a significant reduction in gelatin degradation and thereby a lower gelatinase activity (decreased number of green dots of cleavage-revealed FITC) in Rac inhibited embryo compared to control (**Fig. 3J**). Since the proteolytic function of macrophages in the AGM is essential to HSPC mobilisation, we assessed the effect of Rac inhibition on haematopoietic organ colonisation. We noticed an increase in HSPCs accumulated in the AGM at 48 hpf (+70±8%, n=6) and consequently a decrease in HSPC accumulated in the CHT at 55 hpf (−41±2%; n=5).

### MMP inhibition affects macrophage shape, behaviour and function

Rac inhibition has an impact on macrophage proteolytic activity and consequently on their function. To assess whether direct inhibition of macrophage proteolytic activity induces a similar behaviour, we soaked embryos in a medium containing SB-3CT MMP inhibitor. We previously demonstrated that ECM degradation occurred as a result of macrophage-secreted MMP-9 around HSPCs to enable their intravasation. We evaluated the direct impact of MMP inhibition on macrophage morphology and noticed a variation in shape distribution: an increase in round shape number and a decrease in star-like and elongated shapes (**Fig. 4A**). Moreover, MMP inhibition affected macrophage migration and behaviour (**Fig. 4B-E, Video 3**). Selected images from **Video 3** displayed a typical example of macrophage migration pattern. Using Colour depth projection we were able to visualise the 3D migration of macrophages in the AGM and noticed that in MMP-9 inhibited embryos, macrophages migrated mainly in 2D. At a given point in time, they adopted a single colour whereas in control embryos we observed dynamic changes indicated by the presence of several colours at one time point (**Fig. 2A-F**). Moreover, **Video 3** showed the macrophages adopted different migration pattern resembling to the leukocyte crawling on vein vessel.

**Figure 4:**
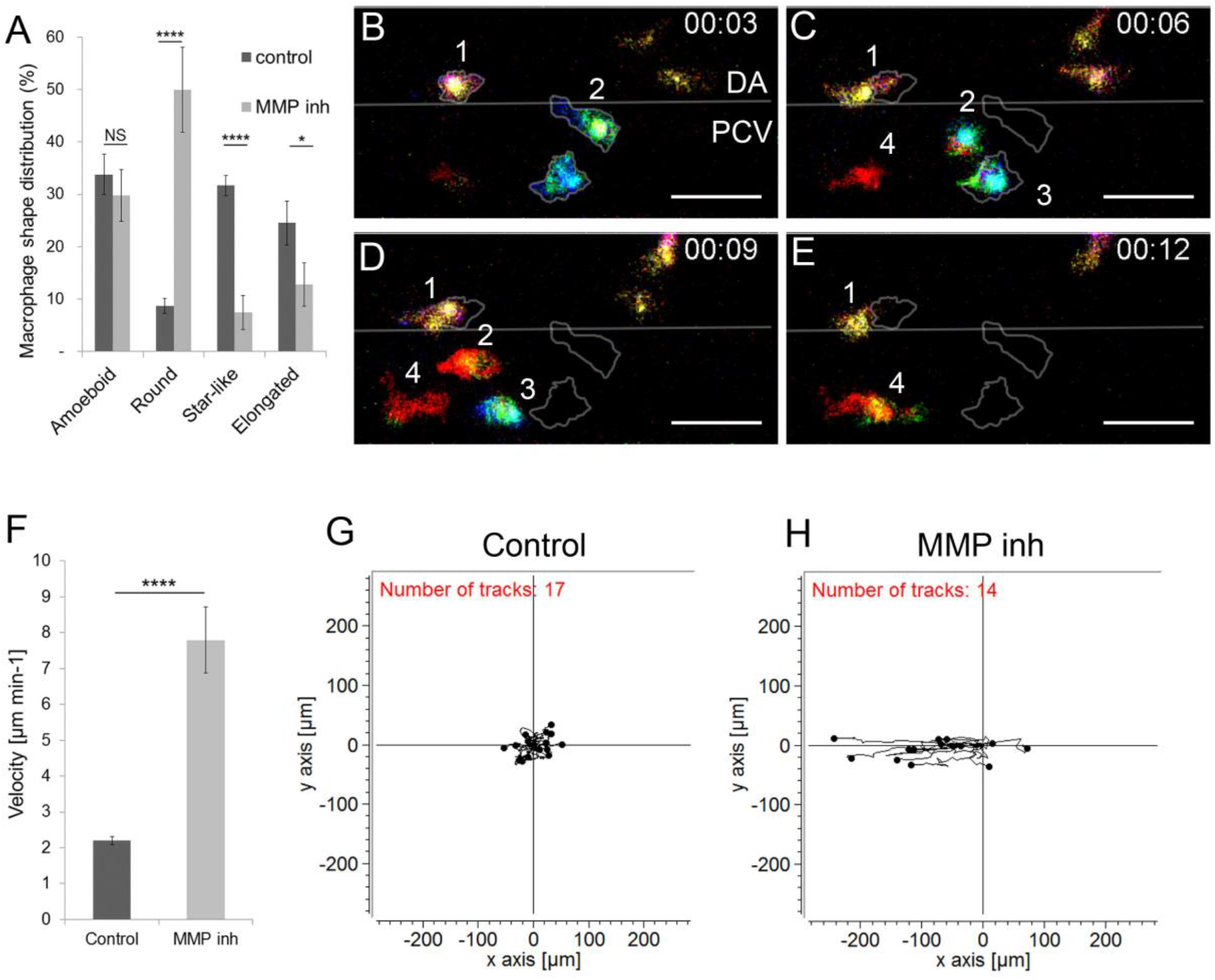
MMP-9 inhibition induces a change in macrophage shape and a transition towards an amoeboid-like migration. (A) Graph compares the macrophage shape distribution in the AGM of MMP-2 and 9 (SB-3CT) - inhibited embryos (MMP inh) to DMSO treated embryos (control) shows an increase in round shape and a decrease in star-like and elongated shapes in MMP inh embryos. N=10 embryos for control and 15 for MMP inh. Data represent the percentage mean for each shape type out of the total number of macrophages in the AGM ± s.e.m. NS = not significant; *p<0.05; ****p<0.0001. (B-E) Selected cropped images from Video 3 displays macrophage shape and migration patterns over time. Numbers point to individual macrophages, time code is expressed in hours and minutes. Grey outlines on panel C-E show the shape and position of macrophages from panel B (3^rd^ minute). (F) Graph showing the velocity of macrophages in control and MMP-inhibited embryos. Data are represented as a mean ± s.e.m., n= 17 macrophages for control and 14 for MMP inhibitor from 4 different embryos, ****p<0.0001. (G-H) Tracking plot diagram represents the migration path and distance of macrophages in the AGM in control and MMP-inhibited embryos measured every minute over 60 minutes. Scale in μm, n=17 macrophages for control and 14 for MMP inhibitor from 4 different embryos. See also Video 3. Scale bar 30 μm. DA, dorsal aorta; PCV, posterior cardinal vein.

Furthermore, we observed that MMP inhibition affected macrophage velocity and directionality. The speed of migration increased more than 3 times compared to the control (from 2.20 ± 0.11 μm.min^−1^ to 7.80 ± 0.92 μm.min^−1^; **Fig. 4F**). Finally, a tracking plot diagram which illustrated the migration path and distance of macrophages in the AGM in control and MMP-inhibited embryos (**Fig. 4G** and **H**) revealed that migration directionality increased from 0.27 to 0.66. Cell tracking showed that macrophages migrated along the vein, in the same direction as the blood flow. We concluded that, MMP inhibition affected both macrophage shape and migration patterns. They adopted a MMP independent migration pattern with increased velocity which was reminiscent of an amoeboid type of migration.

## Discussion

In this study we characterised in zebrafish embryos the macrophage population present in the AGM with a known proteolytic function (Travnickova et al., 2015). We reported the existence of four macrophage morphological subgroups. Previous studies performed *in vitro* described two major morphological types, elongated and rounded shapes (McWhorter et al., 2013). Using *in vivo* analyses we were able to identify two additional morphological shapes: amoeboid and star-like shapes. *In vivo* observations revealed the presence of a higher number of macrophage subgroups in contrast to conclusions drawn from assays on 3D matrices, thereby suggesting the importance of *in vivo* modelling to complete results obtained *in vitro*. Using high resolution live imaging in conjunction with macrophage shape descriptor analysis we devised a novel tool that enabled to quantify *in vivo* the dynamics and morphological plasticity of macrophages.

While macrophages were thought to exclusively migrate using an amoeboid mode (Friedl and Weigelin, 2008), Dr Parini’s group demonstrated their capacity to also use a mesenchymal migration mode (Cougoule et al., 2012). In line with this last study, we describe the mesenchymal macrophage migration process *in vivo* in zebrafish embryos. Macrophages revealed an increase in shape plasticity which confirmed the outcome of previous studies performed *in vitro* (Cougoule et al., 2012).

Previous studies highlighted the significance of the role played by Rac signalling in cytoskeleton organisation during the mesenchymal migration of cells (Sanz-Moreno and Marshall, 2010). Our study performed *in vivo* during the establishment of haematopoiesis in zebrafish embryos also demonstrated that the mesenchymal migration of macrophages was Rac signalling dependent. Going further, we observed that Rac signalling inhibition affected not only macrophage migration but also their proteolytic function and their phenotype. Indeed, upon Rac inhibition macrophages lose their ability to degrade the ECM matrix. We also observed that this treatment significantly reduced macrophage velocity and morphological plasticity. Moreover, live imaging revealed that macrophages develop and keep longer membrane extensions and that they remained longer in a specific location. Our study confirmed previous *in vitro* observations showing that Rac1-/- macrophages cultured on plastic exhibited additional membrane extensions when compared to control macrophages (Wheeler et al., 2006).

We finally observed that the inhibition of the macrophage proteolytic function induces their transition into a different type of migration mode corresponding to the adaptation of macrophages to their new environment. They adopted a round shape with an amoeboid migration. Macrophages were no longer able to migrate within the AGM stroma and they moved along the vein wall.

Proteolytic macrophages in the AGM exhibited a high functional similarity to macrophages found in solid tumours referred to as tumour associated macrophages (TAM). TAM play a significant part in ECM remodelling through proteinase releases (mainly MMP-2 and 9) and allow tumour cells to join the bloodstream and to seed in secondary sites (Condeelis and Pollard, 2006). Therefore, a current strategy is to target TAM to combat cancer (Panni et al., 2013). Several approaches based on macrophage depletion (clodronate liposomes) or functional modification (broad spectrum MMP inhibitors) did not succeed and failed during clinical trials due to low specificity and the amount of side effects (Panni et al., 2013; Turk, 2006). Expanding our knowledge from a purely molecular standpoint toward an in-depth understanding of behaviour and requirements in migration and site infiltration using adapted *in vivo* models, would complement existing studies and enable us to develop more targeted immunotherapeutic solutions.

## Materials and Methods

### Zebrafish husbandry

Wild-type and transgenic lines were maintained in compliance with the Institutional Animal Care and Use protocols. The following transgenic lines were used in this study: *Tg(Mpeg1:mCherry-F)* (Ellett et al., 2011; Travnickova et al., 2015) for macrophage membrane marking and *Tg(kdrl:eGFP)* (Beis et al., 2005) for vessel endothelium labelling. Embryos were kept in the presence of 1-phenyl-2-thiourea to prevent melanin pigmentation (Westerfield, 2000) and staged as described by Kimmel et al (Kimmel et al., 1995). All experiments were performed in accordance with the protocol CEEA-LR-13007 approved by the Animal Care and Use Languedoc-Roussillon Committee.

### Live Imaging

Zebrafish embryos (lateral views, rostral to the left) were embedded in 0.7% low melting agarose and imaged using a Zeiss LSM510 confocal microscope through a 40x water immersion objective with a 1024×256 pixel resolution at 28°C. All live imaging experiments were performed at 46-48 hpf and all time-lapse imaging occurred at an acquisition rate of one minute at a 1μm z-interval. The acquisitions were performed using ZEN2009. Image processing such as maximum intensity projections, 3D view, and overall image contrast adjustment were performed using Fiji software.

### Inhibitor treatment

Embryos were soaked in MMP-2 and 9 inhibitors SB-3CT (Enzo Life Sciences) 9 μM or NSC23766 Rac inhibitor (Tocris) 50 μM or DMSO 0.25% as a control from 5-prim stage (25 hpf) to 46-48 hpf. For stock solution, inhibitors were dissolved in DMSO at a 10mM concentration and stored at −20°C.

### Image processing and macrophage shape analysis

Confocal stacks of membrane-labelled macrophages were projected using a maximum intensity projection and 2D images were binarised using an automatic threshold. The following shape descriptors were evaluated using the Fiji plugin Particle analysis: area (μm^2^), perimeter (μm), circularity and roundness. The elongation factor was manually measured by dividing the longest axis of the object by its longest perpendicular axis (x/y). Objects with an area under 80 μm^2^ were excluded from the further analysis. Circularity was calculated using the following formula: 4π x (area/perimeter^2^). This parameter varied from 0 (linear polygon) to 1 (perfect circle). Circularity was used to set apart round objects (circularity > 0.2) and roundness and elongation factor enabled us to break down non-round subjects into 3 subgroups: elongated, amoeboid and star-like shaped. Roundness was calculated using the following formula: 4 x {area/ [π x (major axis)^2^]} and varied from 0 (linear polygon) to 1 (perfect round). Supplementary table 1 shows the mean values of circularity, roundness and elongation factor measured for each of the above listed subgroups.

### Cell tracking and velocity measurement

Maximum intensity projections of 60 minute time-lapses acquired every minute were analysed using a manual tracking plugin in Fiji. Measured data were transferred into a Chemotaxis and Migration tool programme (Ibidi) to design tracking and rose plots (Figure 2G-H for rose plots, 3G-H and 4G-H for tracking plots). A rose diagram maps single counts of the position of every macrophage in a selected area (black and grey sectors of angle π/18) every minute over 60 minutes with an (x,y 0,0) starting point. The tracking plot diagram represents the migration path and distance of macrophages in the AGM with an x,y 0,0 starting point, being measured every minute over 60 minutes. The average of single macrophage velocities (μm min^−1^) during 15-60 minutes were used for analysis. The evaluation of the directionality was performed using a Rayleigh statistical test for the uniformity of a circular distribution of points (end points of single macrophages). All analyses were conducted using the Chemotaxis and Migration tool software (Ibidi).

### Fin amputation for oriented migration analysis

Caudal fin amputation was performed with a sterile scalpel at 44 hpf, posterior to muscle and notochord under anaesthesia with 0.016% Tricaine (ethyl 3-aminobenzoate, Sigma Aldrich). 4 h post amputation embryos were mounted and imaged as described above.

### *In vivo* zymography

The *In vivo* zymography was performed according to Crawford’s protocol (Crawford and Pilgrim, 2005). A working solution, 1 mg ml^−1^ of fluoresceinated gelatin (Gelatin-FITC,Anaspec) in PBS was injected (4-5 ng) into muscles between 4^th^ and 5^th^ somite at 42 hpf. Imaging was performed following the injections. Embryos were incubated in DMSO or Rac inhibitor from 25 hpf up to the Gelatin-FITC injections.

### Statistical analysis

Normal distributions were analysed using the Shapiro-Wilk test. Non-Gaussian data were analysed using the Wilcoxon test, Gaussian with Student t-test. P<0.05 was considered as statistically significant (symbols: ****p<0.0001 *** p<0.001; ** p<0.01; * p<0.05) Statistical analyses were performed using the R software.

## Acknowledgements

We would like to thank E. Lelièvre, M. Rossel and G. Lutfalla for their critical review of this manuscript; V. Diakou and the MRI facility for their assistance, A. Sahuquet for his help in the particle analysis. This work was supported by grants from the ARC, Chercheur d’Avenir - Région Languedoc-Roussillon, FRM and ATIP-Avenir. J.T. was supported by a fellowship from the MESR and FRM (FDT20150532507), M.N.-C. by a fellowship from the Université de Montpellier.

## Author contributions

J.T., and K.K. designed the project and the experiments, J.T., S.N., M.N.-C and N.A. performed the experiments and analysed the results. J.T. and K.K. wrote the manuscript with the input of S.N., M.N.-C. and F.D.

## Ethics

All animal experiments described in the present study were conducted at the University of Montpellier in compliance with European Union guidelines for handling of laboratory animals (http://ec.europa.eu/environment/chemicals/lab_animals/home_en.htm) and were approved by the Direction Sanitaire et Vétérinaire de l’Hérault and Comité d’Ethique pour l’Experimentation Animale under reference CEEA-LR-13007.

## Abbreviations

AGM: Aorta-Gonad-Mesonephros
HSPC: Haematopoietic stem and progenitor cells
MMP: Matrix metalloproteinases
TAM: Tumour associated macrophages

---

**Supplementary table 1:** Measured average values (± s.e.m.) of shape descriptors for each shape subgroup. N = 20 embryos.

| **Shape** | **Circularity** | **Roundness** | **Elongation factor** |
| --- | --- | --- | --- |
| **Round** | $0.19 \pm 0.01$ | $0.65 \pm 0.04$ | $1.55 \pm 0.10$ |
| **Amoeboid** | $0.07 \pm 0.004$ | $0.49 \pm 0.01$ | $2.04 \pm 0.05$ |
| **Star-like** | $0.07 \pm 0.004$ | $0.70 \pm 0.01$ | $1.56 \pm 0.07$ |
| **Elongated** | $0.07 \pm 0.005$ | $0.35 \pm 0.01$ | $3.45 \pm 0.14$ |

**Video 1:** Macrophage 3D migration in the AGM during haematopoiesis.

Representative time-lapse colour-depth projections of *Tg(mpeg1:mCherry)* embryo at 46 hpf illustrate macrophages migration occurring in a non-directional manner and through different depth by appropriately changing colour. Image stacks were acquired every minute over 60 minutes at a 1 μm interval with 1024×256 pixel resolution using the LSM510 Zeiss confocal microscope equipped with a 40x water immersion objective. Scale bar 30 μm, time code in hours and minutes.

**Video 2:** The migratory behaviour of macrophage changes after Rac inhibition.

Combined representative time-lapse colour-depth projections of *Tg(mpeg1:mCherry)* embryos at 46 hpf draws a comparison between macrophage migration in DMSO-treated (control, top) embryos and that of Rac-inhibitor (Rac inh, bottom) treated embryos. Rac-inhibited macrophages display slower migration modes. They change shapes and migration direction less often, and form very long membrane extensions. Image stacks were acquired every minute over 60 minutes at a 1 μm interval with 1024×256 pixel resolution using the LSM510 Zeiss confocal microscope equipped with a 40x water immersion objective. Scale bar 30 μm, time code in hours and minutes.

**Video 3:** MMP inhibition induces mesenchymal-amoeboid transition of macrophage migration. Combined representative time-lapse colour-depth projections of *Tg(mpeg1:mCherry)* embryos at 46 hpf draw a comparison between macrophage migration in DMSO-treated (control, top) and MMP-2 and 9 inhibitor (MMP inh, bottom) treated embryos. MMP-inhibited macrophages migrate faster, adopt a round shape, change the depth of their displacement less often and migrate partially inside the bloodstream. Image stacks were acquired every minute over 60 minutes at 1 μm interval with 1024×256 pixel resolution using the LSM510 Zeiss confocal microscope equipped with a 40x water immersion objective. Scale bar 30 μm, time code expressed in hours and minutes.

